# Fructooligosaccharide Supplementation Improves Glucose Homeostasis in Human-Relevant hyperglycemic Diet-Induced Obese Mice

**DOI:** 10.64898/2026.06.23.733678

**Authors:** Uday Saxena, Shreya Shahapur, Sadiya Mehboob, Priyanka Jadhav, Tanisha Samal, Gopi Kadiyala, Markendeya Gorantla

## Abstract

Fructooligosaccharides (FOS) are prebiotic fibers that influence gut microbiota and host metabolic function. In a diet-induced obesity (DIO) mouse study, FOS supplementation was compared with PBS-treated obese controls. Blood glucose was markedly lower at Day 42 (221.9 ± 7.8 vs 138.3 ± 9.0 mg/dL), and remained lower at Day 56. FOS reduced body-weight gain from 8.4 ± 0.9 g in PBS controls to 2.6 ± 0.2 g, corresponding to an approximate 69.5% reduction in gain over Days 1-70. Cumulative feed consumption was not significantly different between PBS and FOS cages, suggesting that the observed metabolic effects were not explained simply by reduced food intake. These data support our thesis that FOS works as an active metabolic ingredient acting through the gut-liver-metabolic axis. Thus, in the present study, dietary FOS supplementation produced marked improvements in glucose homeostasis in a severe DIO model characterized by diabetic-range hyperglycemia that more closely resembles poorly controlled human type 2 diabetes.

**HIGHLIGHTS:**

- Fructooligosaccharide (FOS) normalized glucose levels in a severe DIO model that mimics poorly controlled human type 2 diabetes.
- Day-42 blood glucose was reduced by ∼37.7% in FOS-treated DIO mice.
- FOS reduced body-weight gain by ∼69.5% versus controls over 70 days.
- Metabolic benefits occurred without a statistically significant reduction in feed intake.
- Findings support a gut–liver–metabolic mechanism rather than simple caloric restriction.
- Data position FOS as an active metabolic ingredient with potential utility in diabetes and metabolic health.

## Introduction

Obesity, insulin resistance, visceral adiposity, and fatty liver disease are interrelated manifestations of metabolic dysfunction. Diet-induced obesity mouse models are widely used to assess interventions that modulate body-weight gain, glucose handling, adiposity, and liver pathology. FOS is a fermentable prebiotic fiber that can alter gut microbial composition and increase gut short-chain fatty acid production. Prior studies have reported that FOS improves glucose homeostasis, visceral adiposity, and hepatic steatosis in obese or high-fat-diet-fed animals. The present analysis supports the idea that FOS actively modulates glucose metabolism. Furthermore, its suggest that the regulation is beyond just limiting food intake as was described in previous studies.

Multiple studies have demonstrated beneficial effects of FOS on glucose tolerance, insulin sensitivity, hepatic lipid metabolism, and body weight regulation in rodent models of obesity and diabetes. These effects are believed to be mediated primarily by involving alterations in microbial fermentation and inflammatory signaling. However, the magnitude of glucose lowering reported in the literature has generally been modest, and the efficacy of higher-dose FOS in severe diabetic-like obesity models remains incompletely characterized.

The present study therefore sought to evaluate the metabolic effects of dietary FOS in a severe DIO model exhibiting marked hyperglycemia, thereby providing an opportunity to assess the therapeutic potential of prebiotic intervention under conditions that more closely resemble advanced human type 2 diabetes. Given the severity of metabolic dysfunction in this model, the study also offered an opportunity to determine whether higher-dose FOS could achieve meaningful glucose normalization beyond the improvements typically reported in milder obesity and prediabetes models.

## Methods

### Animal Study Design

Male BALB/c mice (6–8 weeks old; n = 7 per group)after one week of acclimatation were utilized in a diet-induced obesity (DIO) study conducted over a 70-day period. Animals were maintained on a high-fat diet (HFD; typically 45–60% kcal from fat) throughout the study to induce obesity and metabolic dysfunction. During the induction phase, mice received fructose-supplemented drinking solution in addition to the HFD. Animals were monitored daily for general health, activity, and tolerability.

### Study Design

The present analysis focused on fructooligosaccharide (FOS)-treated animals and PBS-treated controls. Animals were allocated to treatment groups according to the original study protocol and maintained on HFD throughout the experimental period. Retroorbital blood samples were collected at baseline and predefined follow-up time points. Primary endpoints included fasting blood glucose, body weight, and cumulative glucose exposure. Secondary endpoints included feed consumption and overall metabolic progression during the study period.

### FOS Administration

Food-grade fructooligosaccharides (FOS) obtained from a commercial supplier in Hyderabad, India, were administered orally once daily at 10% (w/v) in phosphate-buffered saline (PBS). Control animals received PBS vehicle alone. Treatment was continued throughout the study period.

### Body Weight Assessment

Individual body weights were recorded longitudinally throughout the study. Mean body weight and body-weight gain were calculated for each group. Total body-weight gain was determined as the difference between Day 70 and Day 1 body weight.

### Fasting Blood Glucose Measurements

Animals were fasted prior to blood collection. Fasting blood glucose concentrations were measured at baseline and predefined study time points using an Abbott glucometer. Longitudinal glucose trajectories were generated for each treatment group and used to assess the progression of diet-induced hyperglycemia and the effects of FOS supplementation on glucose homeostasis.

### Feed Consumption Analysis

Feed consumption was recorded at the cage level and normalized to estimated grams consumed per mouse per day. Seven-day rolling averages were generated to visualize temporal trends in food intake. Cumulative feed consumption over the study period was also calculated.

### Glucose Area-Under-the-Curve (AUC) Analysis

Cumulative glucose exposure was quantified by calculating glucose area-under-the-curve (AUC) from Day 0 through Day 56 using the trapezoidal rule. Individual animal fasting blood glucose measurements were used for all AUC calculations. AUC values were utilized as an integrated measure of glycemic burden throughout the study period.

### Statistical Analysis

Data are presented as mean ± standard error of the mean (SEM). Comparisons between PBS and FOS groups were performed using two-sided Welch’s t-tests. Statistical significance was defined as p < 0.05. Graphical analyses were generated using group means and SEM values.

### Ethics Statement

All animal procedures were conducted in accordance with Institutional Animal Ethics Committee (IAEC) requirements and complied with CPCSEA guidelines for the care and use of laboratory animals.

## Results

### FOS reduced body-weight gain relative to PBS control

PBS control animals increased body weight on the high fat diet from 26.7 ± 2.0 g on Day 1 to 35.1 ± 1.8 g on Day 70. FOS-treated animals increased from 24.4 ± 0.3 g to 27.0 ± 0.3 g over the same period. The resulting body-weight gain was 8.4 ± 0.9 g in PBS controls versus 2.6 ± 0.2 g in FOS animals (Welch t-test p=0.0005).

**Figure 1.**
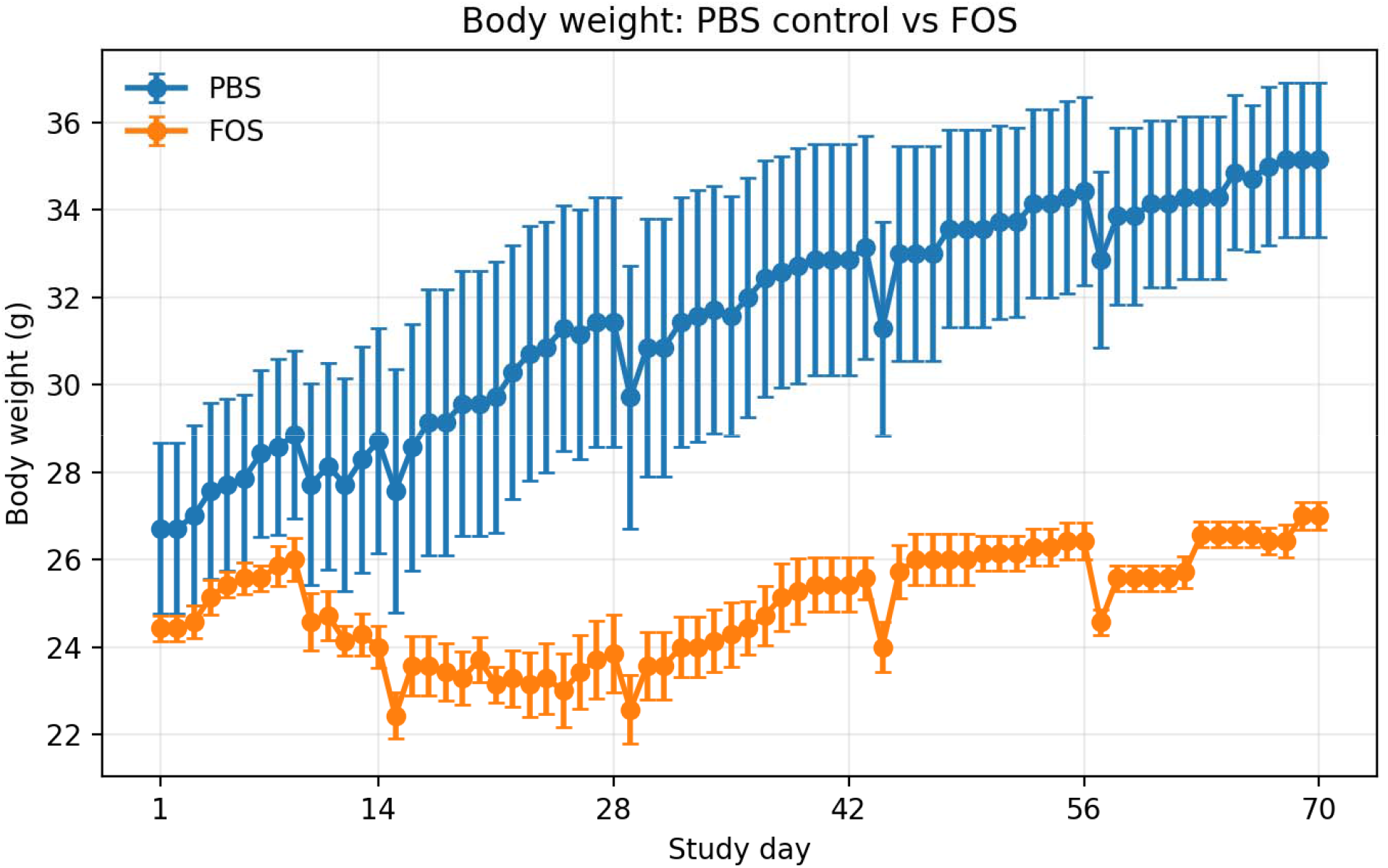
Body-weight trajectory in PBS control and FOS-treated DIO mice. Data are mean ± SEM, n=7/group.

### FOS improved blood-glucose control

At Day 28, mean blood glucose was 238.0 ± 7.7 mg/dL in PBS controls versus 135.1 ± 12.0 mg/dL in FOS-treated mice. At Day 42, glucose was 221.9 ± 7.8 mg/dL in PBS controls versus 138.3 ± 9.0 mg/dL in FOS animals (approximately 37.7% lower; p=0.00002). At Day 56, FOS glucose values remained lower than PBS controls (196.3 ± 6.5 for FOS vs 219.6 ± 5.2 mg/dL; for PBS vehicle group=0.0173).

**Figure 2.**
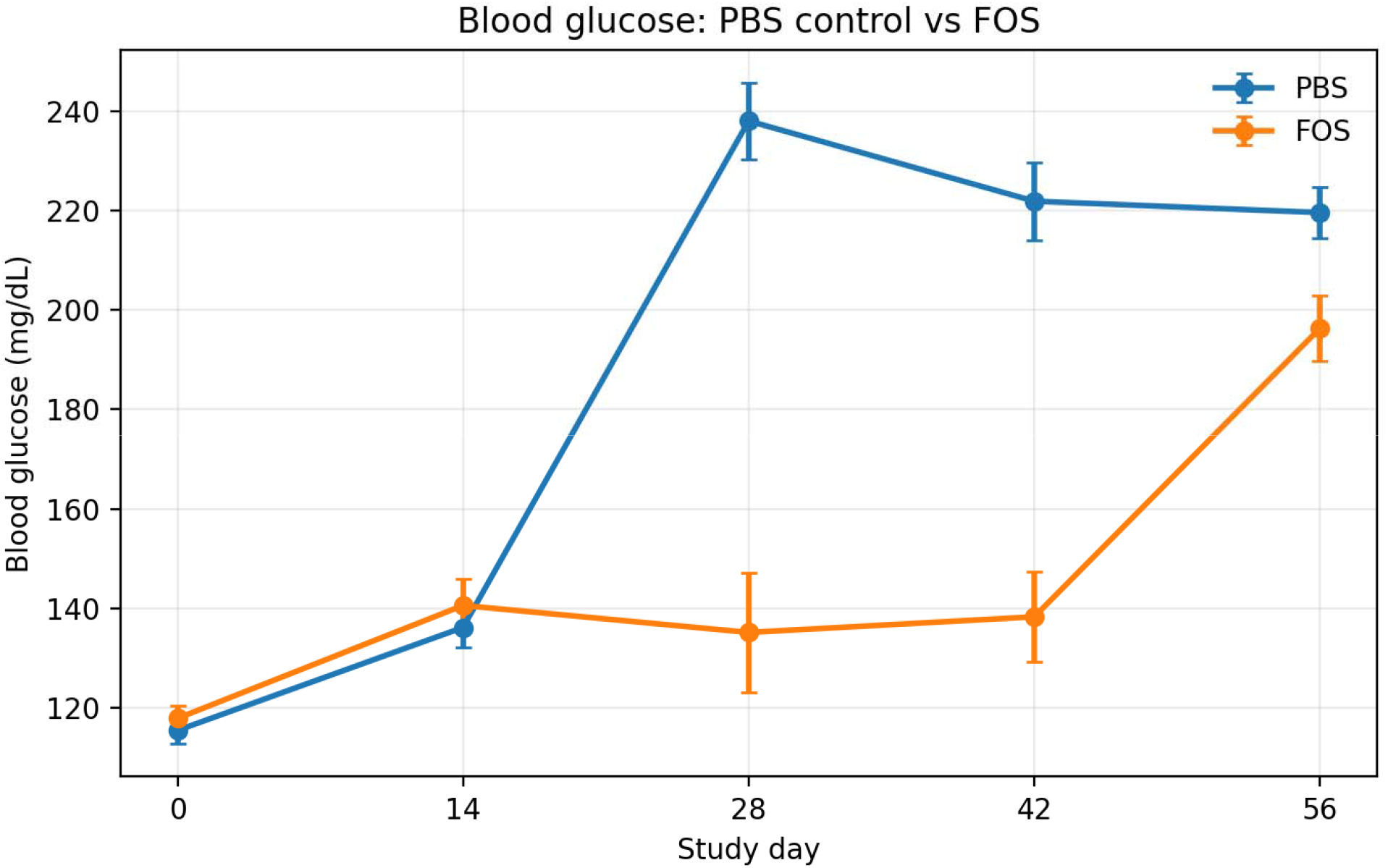
Blood-glucose trajectory in PBS control and FOS-treated DIO mice. Data are mean ± SEM, n=7/group.

### Figure 3. Glucose AUC Analysis

FOS supplementation produced a marked improvement in overall glycemic control as assessed by glucose area under the curve (AUC) analysis also. Compared with the disease control group, FOS-treated animals exhibited a substantially lower glucose AUC, indicating reduced cumulative glycemic exposure throughout the study period. The reduction in AUC was consistent with the lower fasting blood glucose values observed in FOS-treated mice, particularly at later time points when hyperglycemia progressively worsened in control animals. While disease control mice demonstrated a continual rise in blood glucose levels, FOS supplementation attenuated this deterioration and maintained glucose concentrations closer to baseline values. These findings suggest that dietary FOS improves glucose homeostasis in DIO mice and delays the progression of obesity-associated hyperglycemia. The magnitude of the effect was notable given that FOS is a non-pharmacological dietary intervention, supporting its potential role as a microbiome-mediated metabolic modulato. Overall, the AUC analysis demonstrates that chronic FOS supplementation significantly reduces cumulative glycemic burden in this human-like model of obesity-associated metabolic dysfunction.

**Figure 3.**
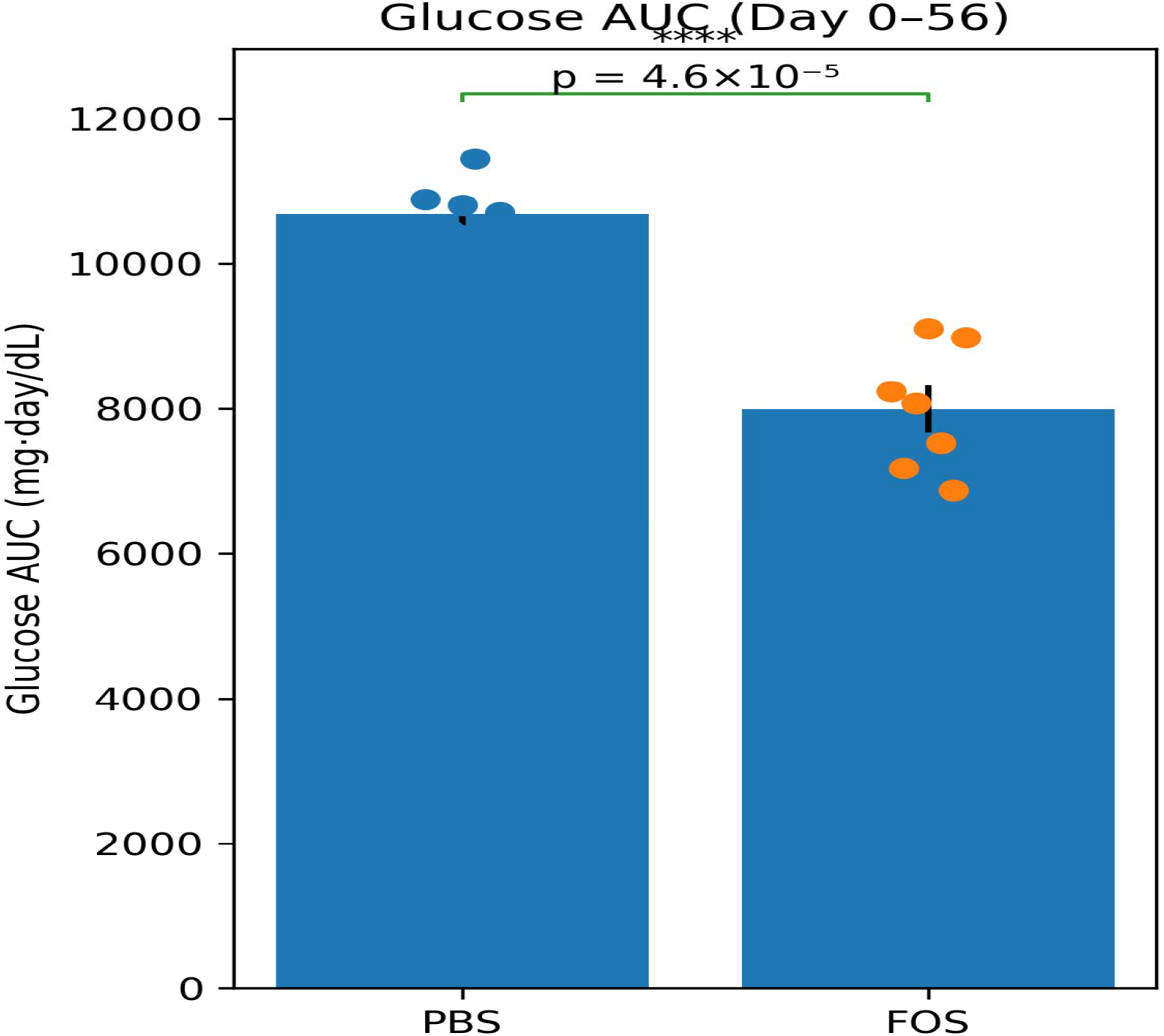
Glucose area-under-the-curve (AUC) analysis in PBS and FOS-treated DIO mice. Individual animal blood-glucose measurements obtained from Day 0 through Day 56 were integrated using the trapezoidal rule. FOS treatment significantly reduced cumulative glucose exposure compared with PBS controls. Mean glucose AUC values were 10,689 ± 159 mg·day/dL for PBS-treated animals and 7,995 ± 325 mg·day/dL for FOS-treated animals, representing a 25.2% reduction in cumulative glucose burden (Welch’s t-test, p=0.000046; n = 7/group).

### Feed consumption does not fully explain the metabolic response

Cumulative feed consumption from Day 1 to Day 70 was 276.3 ± 58.0 g/mouse in PBS cages and 234.1 ± 17.6 g/mouse in FOS cages (cage-level analysis, n=2 cages/group; p=0.598). Although the FOS group trended lower, the cage-level comparison was not statistically significant, supporting the interpretation that improved glucose and reduced weight gain were not driven solely by reduced food availability or intake.

**Figure 4.**
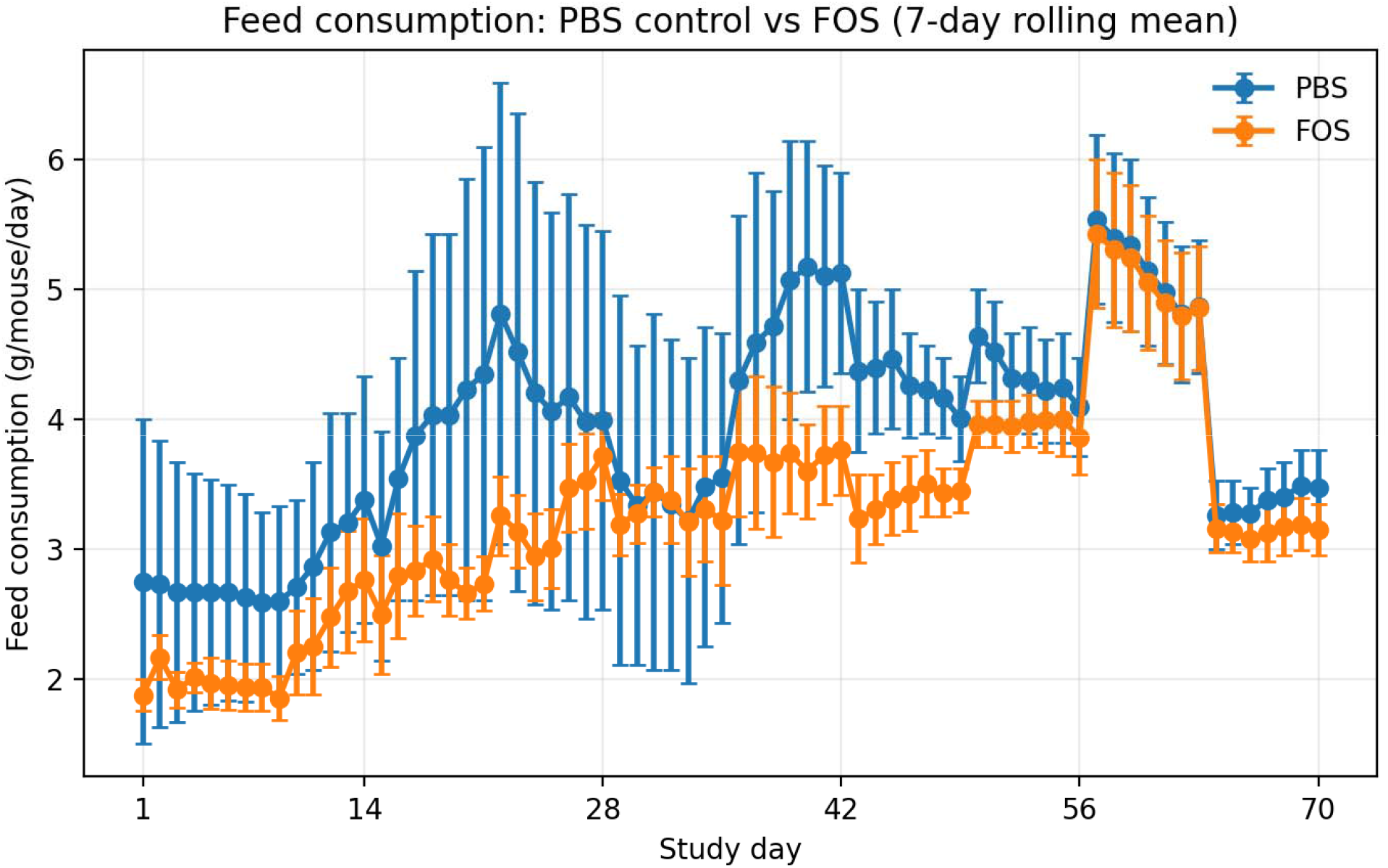
Feed consumption normalized per mouse and shown as a 7-day rolling mean. Data are cage-level mean ± SEM, n=2 cages/group.

### Summary of FOS effects

**Figure 5.**
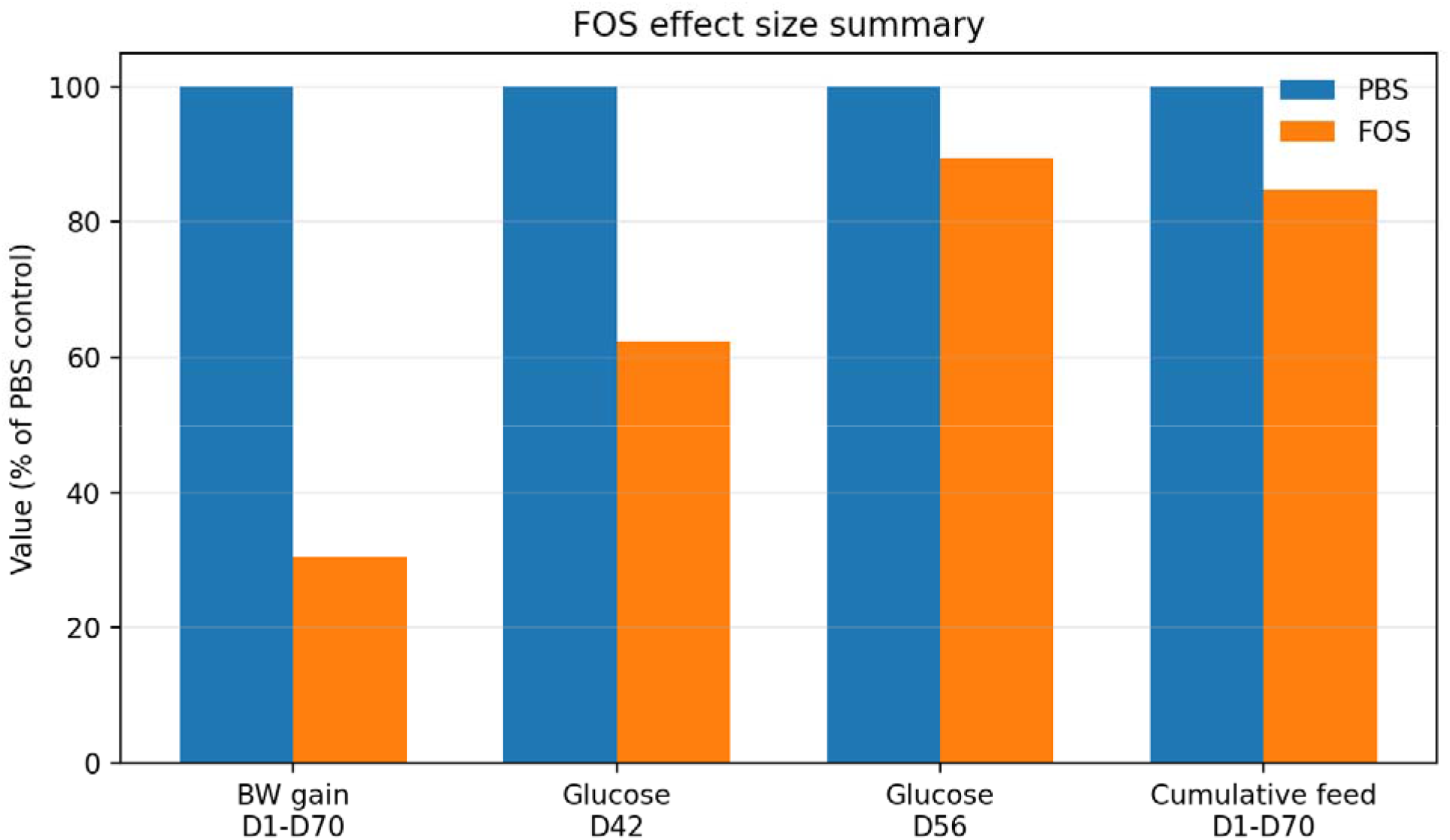
Normalized effect summary for all parameters recorded. PBS control is set to 100% for each endpoint.

### Quantitative Summary Table

| Endpoint | PBS control | FOS | Interpretation |
| --- | --- | --- | --- |
| Body weight Day 1 | 26.7 ± 2.0 g (n=7) | 24.4 ± 0.3 g (n=7) | FOS animals began slightly lighter. |
| Body weight Day 70 | 35.1 ± 1.8 g (n=7) | 27.0 ± 0.3 g (n=7) | FOS group remained substantially lower. |
| Body-weight gain Day 1-70 | 8.4 ± 0.9 g (n=7) | 2.6 ± 0.2 g (n=7) | ~69.5% lower gain; p=0.0005. |
| Blood glucose Day 42 | 221.9 ± 7.8 mg/dL (n=7) | 138.3 ± 9.0 mg/dL (n=7) | ~37.7% lower; p=0.00002. |
| Blood glucose Day 56 | 219.6 ± 5.2 mg/dL (n=7) | 196.3 ± 6.5 mg/dL (n=7) | ~10.6% lower; p=0.0173. |
| Cumulative feed Day 1-70 | 276.3 ± 58.0 g/mouse | 234.1 ± 17.6 g/mouse | Cage-level trend lower but not significant; p=0.598. |

## Discussion

In the present DIO study, FOS supplementation produced a clear metabolic benefit, including reduced body-weight gain, improved blood glucose, and a non-significant trend toward lower feed consumption. The pattern of response suggests that FOS acts primarily as a gut-liver-metabolic modulator rather than as a simple appetite suppressant. The robust metabolic effects observed in the present study were achieved using a dietary FOS concentration on the higher side to that employed in several published rodent studies (5– 13%).

Our results are broadly consistent with prior studies (see Table below) showing that FOS and related fermentable fibers improve metabolic dysfunction in obese rodents. Published models of FOS supplementation shown below, have reported reductions in steatohepatitis, visceral adiposity, chronic inflammation, and high-fat-diet-associated fat accumulation. The current study aligns well with that literature, particularly in the strong glucose effect observed at Day 28 and Day 42.

| **Study** | **Model** | **Duration** | **Key Findings** | **Weight Effect** | **Distinctive Feature** |
| --- | --- | --- | --- | --- | --- |
| Takai et al., 2020 | NASH obese mice | 12–16 weeks | Reduced steatohepatitis, visceral adiposity, inflammation | Moderate | Strong liver-fat effect |
| Beserra et al., 2018 | Meta-analysis | Multiple | Improved glucose and metabolic endpoints | Variable | Broad evidence base |
| Parnell & Reimer, 2009 | Overweight humans | 12 weeks | Gut hormones altered, weight reduction | Modest | Human translational evidence |
| Present Study | DIO mice | 70 days | Marked glucose improvement with limited effect on feed intake | ~69.5% reduction in weight gain | Large glucose effect despite similar feed intake |

A key observation in our studies is that the improvement in glucose control and body-weight gain was not fully explained by feed consumption. Feed intake data showed that cumulative intake was not significantly different between groups. This supports the possibility that FOS improved metabolic quality through mechanisms beyond simple caloric restriction.

Mechanistically, the observed metabolic effects are consistent with modulation of the gut– liver axis. Fructooligosaccharides are not substantially absorbed in the upper gastrointestinal tract and therefore exert their primary biological activity within the intestinal lumen. Fermentation of FOS by the gut microbiota generates metabolites that are transported directly to the liver through the portal circulation, providing a direct communication pathway between the intestine and hepatic metabolic tissues. Through this gut–liver axis, FOS may influence hepatic glucose production, lipid handling, insulin sensitivity, and systemic energy metabolism. The substantial reduction in glucose exposure observed in the present study, despite the absence of a statistically significant reduction in feed intake, is consistent with improved metabolic regulation rather than simple caloric restriction. The substantial reduction in glucose AUC observed with FOS supplementation may reflect modulation of hepatic glucose homeostasis through the gut–liver axis. Previous studies have demonstrated that FOS can influence pathways regulating gluconeogenesis, insulin sensitivity, and incretin signaling, including reductions in PEPCK and G6Pase expression and enhancement of GLP-1-mediated glucose control. A human metabolic study found that **20 g/day FOS reduced basal hepatic glucose production**, which directly supports the gut–liver axis hypothesis.

Compared with the published FOS literature, the novelty of the present data is not just the basic observation that FOS improves DIO-associated metabolic dysfunction. Rather, the differentiating feature is the quantitative pattern: marked suppression of weight gain and strong glucose improvement with no statistically significant decrease in cage-level feed consumption. These findings support the view that FOS should be considered an active metabolic ingredient rather than merely a sweetener, bulking agent, or formulation excipient.

The magnitude of glucose lowering observed in the FOS-treated group was greater than typically reported in published diet-induced obesity (DIO) studies. Several factors may explain this finding. First, the disease-control animals in the present study developed marked hyperglycemia, with glucose values reaching approximately 220–250 mg/dL, representing a more severe metabolic phenotype than that observed in many published DIO models, where glucose concentrations commonly range from 180–220 mg/dL. This provided a larger therapeutic window for intervention and may have amplified the observable treatment effect.

Second, FOS was administered at approximately 10% of the diet, which is at the upper end of doses commonly employed in rodent metabolic studies. Higher dietary FOS exposure is known to enhance microbial fermentation and short-chain fatty acid production, leading to greater effects on host metabolism.

Third, the duration of treatment was sufficient to permit microbiome remodeling and downstream metabolic adaptation. Unlike pharmacological glucose-lowering agents, prebiotic interventions typically require prolonged exposure to exert maximal effects through alterations in gut microbial composition, intestinal barrier function, inflammatory signaling, and hepatic metabolism.

The observed glucose reductions may also have been supported by modest reductions in food intake. However, the concurrent improvements in glucose control, body weight trajectory, and other metabolic endpoints suggest that the effects extend beyond simple caloric restriction. Mechanistically FOS may act primarily through modulation of the gut– liver axis, resulting in improved hepatic insulin sensitivity as well as reduced hepatic glucose production.

Importantly, the glucose concentrations observed in the disease-control animals are more consistent with a diabetic phenotype than with simple obesity-associated insulin resistance. In this regard, the present model may more closely resemble poorly controlled human type 2 diabetes than the milder prediabetic states frequently reported in DIO studies. The ability of FOS treatment to reduce glucose values toward near-normal levels under these conditions suggests a biologically meaningful metabolic effect and supports further investigation of FOS-based interventions in advanced metabolic disease.

### Potential Application of FOS in Prediabetes and Type 2 Diabetes

Prediabetes is characterized by progressive insulin resistance, elevated postprandial glucose excursions, and gradual deterioration of glucose homeostasis. Dietary prebiotics such as fructooligosaccharides, have been shown in both animal and human studies to modulate the gut microbiome, increase production of short-chain fatty acids, improve gut barrier integrity, and enhance secretion of incretin hormones including GLP-1 and PYY. These mechanisms may contribute to improved insulin sensitivity and better glycemic control.

In the current DIO model, FOS-treated animals maintained substantially lower blood glucose levels than untreated controls despite prolonged exposure to a high-fat diet. This observation suggests that FOS may help delay or prevent progression from metabolic dysfunction toward overt diabetes. Such an effect would be particularly valuable in individuals with prediabetes, where safe nutritional interventions capable of preserving metabolic homeostasis are highly desirable.

In established type 2 diabetes, FOS is not expected to replace pharmacologic therapies but may serve as an adjunctive nutritional approach. Potential benefits include improved postprandial glucose control, enhanced satiety, support of beneficial gut microbial populations, and reduction of metabolic inflammation. Because FOS has an established safety profile, translation into functional food, beverage, or nutritional supplement formats is feasible for glucose control.

## Acknowledgements

We sincerely thank the team at Cology Biosciences Pvt Ltd located in Hyderabad, India, for helping us with studies evaluating the effects of FOS in murine models. All in vivo studies were conducted in accordance with the Institutional Animal Ethics Committee (IAEC) and complied with CPSCEA guideline

## References

Gélineau A et al. Fructooligosaccharides benefits on glucose homeostasis upon high-fat diet feeding require type 2 conventional dendritic cells. Nature Communications. 2024;15:5413. This study demonstrated improved glucose tolerance and HOMA-IR in HFD-fed mice receiving FOS despite minimal effects on body weight and food intake.

Takai A. et al. Fructo-oligosaccharides ameliorate steatohepatitis, visceral adiposity and associated chronic inflammation in an obese mouse model of NASH. URL: https://pmc.ncbi.nlm.nih.gov/articles/PMC7045471/

FOS / oligofructose systematic review in animal metabolic studies. URL: https://link.springer.com/article/10.1186/s12986-018-0245-3

Dietary supplementation of fructooligosaccharides and hepatic steatosis. URL: https://ffhdj.com/index.php/ffhd/article/view/130

Delzenne NM, Cani PD. Interaction between obesity and the gut microbiota: relevance in nutrition. Annual Review of Nutrition. 2011;31:15–31.

Cani PD, Neyrinck AM, Fava F, et al. Selective increases of bifidobacteria in gut microflora improve high-fat-diet-induced diabetes in mice through a mechanism associated with endotoxaemia. Diabetologia. 2007;50:2374–2383.

Parnell JA, Reimer RA. Weight loss during oligofructose supplementation is associated with decreased ghrelin and increased peptide YY in overweight and obese adults. American Journal of Clinical Nutrition. 2009;89:1751–1759.

Takai A, Kikuchi K, Ichimura M, et al. Fructo-oligosaccharides ameliorate steatohepatitis, visceral adiposity and associated chronic inflammation in an obese mouse model of NASH. BMC Gastroenterology. 2020;20:16.

Roberfroid MB. Introducing inulin-type fructans. British Journal of Nutrition. 2005;93(Suppl 1):S13–S25.

Slavin J. Fiber and prebiotics: mechanisms and health benefits. Nutrients. 2013;5:1417–1435.

Beserra BTS, Fernandes R, do Rosario VA, et al. A systematic review and meta-analysis of the preclinical effects of fructooligosaccharides on metabolic outcomes. Nutrition Journal. 2018;17:18.

Daubioul CA, Horsmans Y, Lambert P, et al. Effects of oligofructose on glucose and lipid metabolism in patients with nonalcoholic fatty liver disease. European Journal of Clinical Nutrition. 2005;59:723–726.

Canfora EE, Jocken JWE, Blaak EE. Short-chain fatty acids in control of body weight and insulin sensitivity. Nature Reviews Endocrinology. 2015.

Chambers ES, Preston T, Frost G, Morrison DJ. Role of gut-derived short-chain fatty acids in appetite regulation and energy homeostasis. Cell Metabolism. 2018.

Koh A, De Vadder F, Kovatcheva-Datchary P, Backhed F. From dietary fiber to host physiology: short-chain fatty acids as key bacterial metabolites. Cell. 2016.

Everard A, Cani PD. Gut microbiota and GLP-1 signaling in obesity and diabetes. Diabetes & Metabolism. 2014.

